# Sequence-Dependent Conformational Transitions of Disordered Proteins During Condensation

**DOI:** 10.1101/2024.01.11.575294

**Authors:** Jiahui Wang, Dinesh Sundaravadivelu Devarajan, Young C. Kim, Arash Nikoubashman, Jeetain Mittal

## Abstract

Intrinsically disordered proteins (IDPs) can form biomolecular condensates through phase separation. It is recognized that the conformation of IDPs in the dense and dilute phases as well as at the interfaces of condensates can critically impact the resulting properties associated with their functionality. However, a comprehensive understanding of the conformational transitions of IDPs during condensation remains elusive. In this study, we employ a coarse-grained polyampholyte model, comprising an equal number of oppositely charged residues—glutamic acid and lysine—whereby conformations and phase behavior can be readily tuned by altering the protein sequence. By manipulating the sequence patterns from perfectly alternating to block-like, we obtain chains with ideal-like conformations to semi-compact structures in the dilute phase, while in the dense phase, the chain conformation is approximately that of an ideal chain, irrespective of the protein sequence. By performing simulations at different concentrations, we find that the chains assemble from the dilute phase through small oligomeric clusters to the dense phase, accompanied by a gradual swelling of the individual chains. We further demonstrate that these findings are applicable to several naturally occurring proteins involved in the formation of biological condensates. Concurrently, we delve deeper into the chain conformations within the condensate, revealing that chains at the interface show a strong sequence dependence, but remain more collapsed than those in the bulk-like dense phase. This study addresses critical gaps in our knowledge of IDP conformations within condensates as a function of protein sequence.

**TOC graphics:** 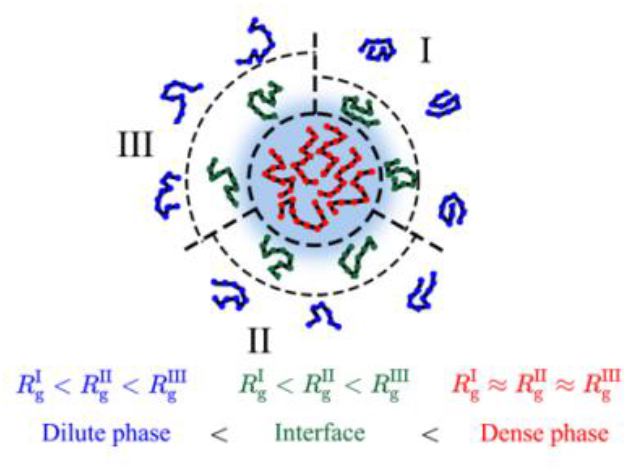

---

Liquid-liquid phase separation (LLPS) is crucial for the formation of biomolecular condensates, which in turn play an important role in vital cellular mechanisms such as gene expression, signal transduction, stress response, and the assembly of macromolecular complexes^1–5^. Intrinsically disordered proteins (IDPs) and disordered regions within proteins have been identified as a primary driving force for the formation of the condensed phase in many cases. It is recognized that the conformation of IDPs in biomolecular condensates and at their interface can critically impact the resulting functionality. For example, recent studies highlighted that the conformational changes of Dcp2 within P-bodies influence mRNA decapping^6^, and conformational expansion of Tau in condensates promotes fibrillization^7^. These results emphasize the important role of protein conformations in dictating the function of biomolecular condensates. Extensive research has been conducted to study protein conformations in the dilute phase^8,9^, elucidating that the conformation of IDPs can be modulated by factors such as sequence composition^10–15^, charge characteristics^16–19^, sequence pattern^20–24^ and solvent environment^25–28^. Sequences comprised of charged residues exhibit a globule-to-coil transition with an increase in the net charge per residue^17^. Quantitative analyses of charge patterns have demonstrated that enhanced charge segregation in the sequence typically leads to more compact conformations in the dilute phase^20,21^. Conformational changes are also modulated by varying electrostatic interactions due to the surrounding solvent environment. Recently, Reddy *et al*. illustrated that a pH shift from neutral to acidic prompted Prothymosin-α to shift from a random coil to a partially collapsed state^29^. In the dense phase, it has been observed experimentally that α-synuclein shifts towards more “elongated” conformations during LLPS^30^, and Tau K18 exhibits expanded conformations within the droplet phase, in contrast to a compact structural ensemble in the dilute phase^31^. A1-LCD protein and its mutated variants have been shown to adopt more extended conformations in the condensates as well through molecular simulations^32^. Despite these excellent prior studies on the conformations of IDPs in the dense phase, there is a general lack of understanding of how the protein conformations change when transitioning from the dilute to the condensed phase; are they always more expanded in condensates than in the dilute phase? What are the polymer scaling properties of IDPs in the dense phase and at the interface, and how do these depend on the protein sequence and dilute phase conformations?

To answer these questions and to decipher the conformational transitions from the dilute to the dense phase during LLPS, we systematically studied a wide range of polyampholyte sequences, which exist in many naturally occurring IDPs^33,34^. Further, these polyampholyte sequences are ideal model systems, since their conformations, as measured by the radius of gyration (*R*_*g*_), can be readily modulated by altering the charge pattern^20,21,35,36^. In particular, we selected 15 sequences composed of glutamic acid (E) and lysine (K) residues (**Figure S1**). Each E-K variant (EKV) consists of 50 residues with an equal number of E and K residues to keep a zero net charge. The degree of charge segregation of EKVs was quantified by the sequence charge decoration (SCD) parameter^21,37^. To allow for a more straightforward comparison with other IDPs, we normalized the SCD values (nSCD) so that nSCD = 0 for the uniformly charge-patterned sequence and nSCD = 1 for the most charge-segregated diblock sequence (normalization method shown in SI text)^38^. The selected 15 EKVs cover diverse conformations in dilute solutions (single chain) at constant temperature (*T* = 300 K), ranging from an ideal-like chain (EKV1) to a semi-compact chain (EKV15). This selection allowed us to probe the conformational properties within the dense phase starting with different chain configurations in the dilute phase. First, we analyzed the conformations for EKVs in the dense phase and as a single chain (**Figure 1**) by calculating the root mean square radius of gyration (*R*_*g*_) from the average of the trace of the gyration tensor (*G*):

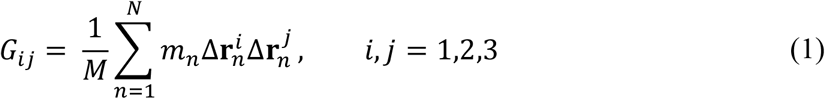

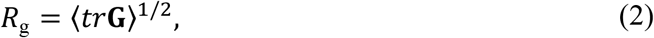

where *M* is the chain’s total mass, *m*_*n*_ is the residue’s mass, and 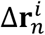 is the vector pointing from the chain’s center of mass to residue *n*, while *i* and *j* are the components in the Cartesian *x, y*, and *z* directions.

**Figure 1.**
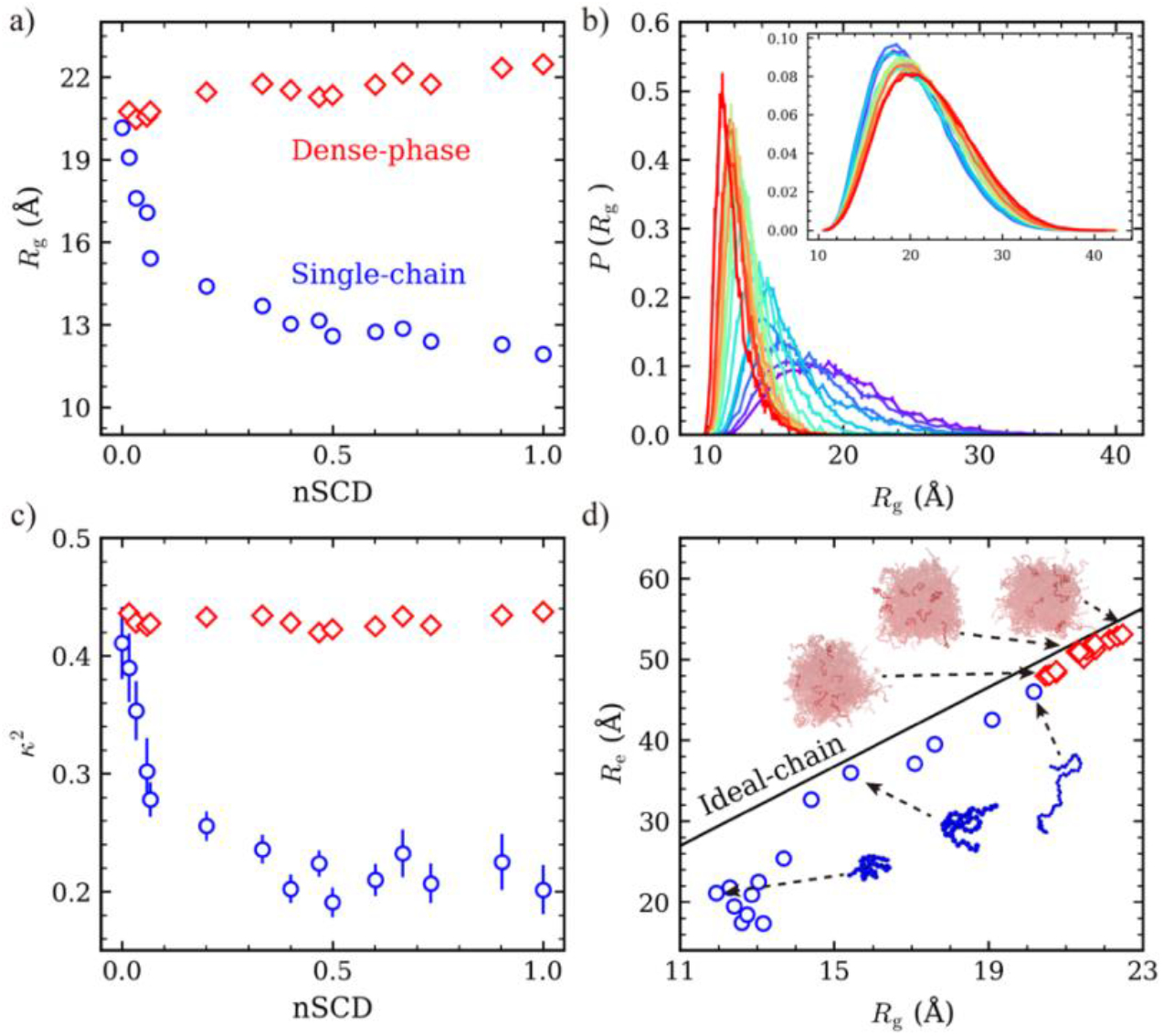
Conformational properties of EKVs in the dense phase (red diamonds) and as a single chain (blue circles). (a) Radius of gyration (*R*_g_) as a function of nSCD. (b) Probability distribution of *R*_g_, *P*(*R*_g_), for EKVs as a single chain. The inset shows *P*(*R*_g_) for EKVs in the dense phase. Line colors, ranging from purple to red, indicate increasing nSCD. (c) Relative shape anisotropy (*κ*^2^) as a function of nSCD. (d) The correlation between end-to-end distance (*R*_e_) and *R*_g_ for the EKVs compared to an ideal chain (solid line). Snapshots show the conformations in the dense phase (red) and as single chains (blue).

For single chains, *R*_*g*_ decreased with increasing nSCD (**Figure 1a**), which is consistent with many previous studies that examined this behavior in detail^20,21,35,39^. To study the conformations in the dense phase, we performed bulk simulations maintaining a constant external pressure (*P* = 0 atm), allowing the systems to adopt their preferred concentration. For EKV1 (nSCD = 0), no dense phase formed at the investigated temperature (*T* = 300 K) because the alternating distribution of positively and negatively charged residues resulted in weak interchain attractions^39^. For all other EKVs, a dense phase formed, where its concentration increased with increasing nSCD (**Figure S2a**). For these cases, as nSCD increased, we observed a corresponding modest rise in *R*_*g*_ within the dense phase, which amounted to an approximate 11.4% increase compared to the single chain *R*_*g*_ of EKV1. Conversely, for a single chain, there was a marked decrease in *R*_*g*_, up to 40.8%, from EKV1 to EKV15 (**Figure 1a**). Consequently, the disparity in *R*_*g*_ between the dense phase and single chains widened as nSCD increased. Specifically, this difference expanded from 8.7% to 88.2% relative to the *R*_*g*_ of single chains, spanning from EKV2 to EKV15. For single chains, the *R*_*g*_ probability distribution, *P*(*R*_*g*_), substantially narrowed with increasing nSCD (**Figure 1b**), which reflects the smaller conformational variety of collapsed blocky EKV sequences. In contrast, *P*(*R*_*g*_) was much broader in the dense phase and slightly broadened with increasing nSCD. These trends indicate that the dense phase of EKVs exhibits a markedly greater diversity of conformations, which depend only weakly on the specific protein sequence, as compared to their single-chain state. These results are consistent with prior experiments of Tau proteins, which also found more expanded conformations and enhanced conformational fluctuations for proteins in droplets^31^.

To characterize the shape of the individual chains, we calculated the average relative shape anisotropy (*κ*^2^) using the three eigenvalues (*λ*_*i*_) of the gyration tensor:

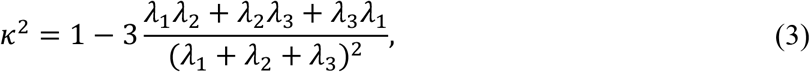

where *κ*^2^ = 0 indicates a spherical configuration, *κ*^2^ ≈ 0.39 an ideal-like chain conformation, and *κ*^2^ = 1 a rod-like structure^40–42^. In the dense phase, *κ*^2^ exhibits only small variations with an increase in nSCD (**Figure 1c**), fluctuating slightly above the value expected for ideal-like chains. In contrast, for a single chain, *κ*^2^ significantly decreases with increasing nSCD, pointing to a coil-to-globule transition. Like the *R*_*g*_ distribution, the probability distribution of *κ*^2^ for single EKVs narrowed when increasing nSCD (**Figure S3**), which suggests restricted conformational variations for sequences with high charge segregation. Furthermore, we compared these conformations with an ideal chain conformation (**Figure 1d**). Interestingly, for EKVs in the dense phase, the end-to-end distance, *R*_e_, closely follows the theoretically expected behavior of an ideal chain, *i*.*e*., *R*_e_ = 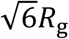 ^43^. However, *R*_e_ of EKVs in single-chain state showed notable deviations from this relationship, particularly pronounced in EKVs with high charge segregation.

These results demonstrate that the conformations observed in the dense phase, as well as those of the sequences with low to moderate degree of charge segregation in the dilute phase, are closely akin to the conformation of an ideal chain. These observed conformational properties can be understood by considering the attractive interactions between monomers. In the dense phase, chains are surrounded by other chains, allowing monomers from a chain to interact with neighboring chains, leading to the observed chain expansion. In contrast, in the dilute phase, monomers can only form intramolecular contacts, thereby leading to collapsed conformation to achieve a state of minimum free energy. Within the dense phase, chain’s conformation nearly mirrors the random-walk characteristics of an ideal chain, which maximizes the conformational entropy (and thus minimizes the free energy)^43^. As a result, in the dense phase, conformations of EKVs exhibited a minor sequence-dependent variation. Conversely, the conformation of an isolated chain in a poor solvent is determined by the intrachain interactions that involve an equilibrium between long-range electrostatic repulsion and attraction, which is substantially modulated by the sequence^20^.

Having established the conformations within the dense and dilute phases, we proceeded to analyze the conformational transitions between these two phases by simulating the EKVs across a series of fixed concentrations, from 0.2 mg/ml to 100 mg/ml. This was achieved by maintaining a constant number of chains (*N* = 500) and modifying the volume of the simulation box (**Figure 2a**). When the concentration exceeds the saturation concentration *c*_*sat*_, some chains spontaneously assemble so that the system phase separates into a dilute and a dense phase. To illustrate this transition in more detail, we chose the sequence EKV5 as an example (**Figure 2a**,**b**), whose *c*_*sat*_ (=0.68 mg/ml) falls within our investigated concentration range. For concentrations *c* ≤ 0.4 m*g*/ml < *c*_*sat*_, the average cluster size remained at one (see SI for technical details of the cluster analysis), indicating that the chains have do not show signs of condensation yet. Correspondingly, the distributions of *R*_*g*_ were identical to the ones measured in our single chain simulations (**Figure S4**).

**Figure 2.**
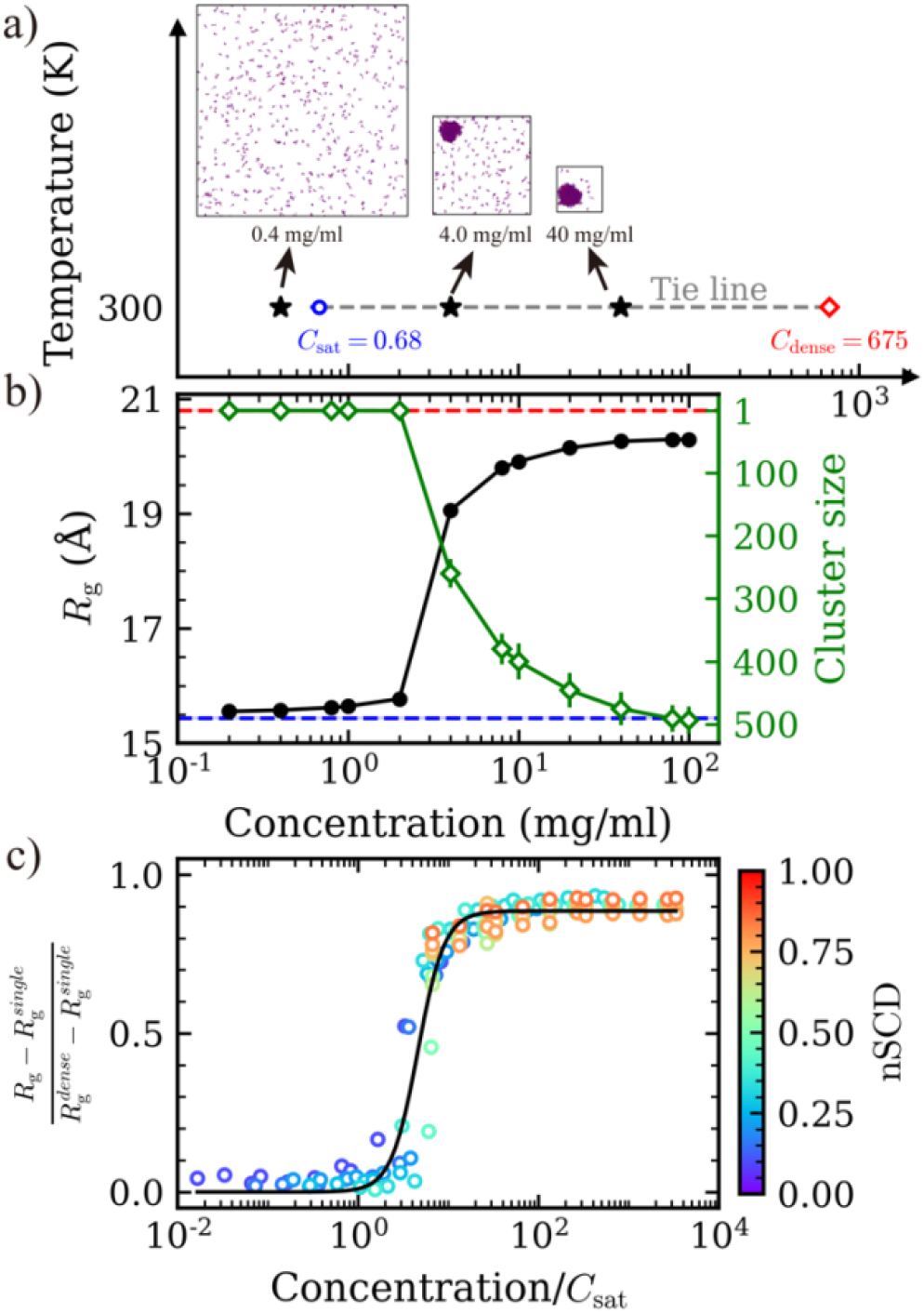
Concentration scan for the EKVs. (a) Tie line for EKV5 at *T* = 300 *K* indicating coexisting dense (*c*_dense_) and dilute (*c*_sat_) phase concentrations along with snapshots at different concentrations. (b) *R*_g_ (black, left *y*-axis) and cluster size (green, right *y*-axis) of EKV5 as functions of concentration. The red and blue dashed horizontal lines represent the *R*_g_ in the bulk dense phase and of a single chain, respectively. (c) Normalized *R*_g_ as a function of normalized concentration for the EKVs. The black solid line is the fitted curve for all simulation data (symbols).

Intriguingly, for *c*_*sat*_ < *c* < 4mg/ml, the average cluster size remains near one, albeit marginally larger, due to the formation of a small number of clusters ranging in size from 2 to 10 assembled chains (**Figure S5a**). The local concentration within these clusters is identical to the dense phase concentrations from our bulk simulations (**Figure S5b**), which suggests that phase separation is indeed occurring above the *c*_*sat*_, even if macroscopic phase separation cannot be observed yet due to system size limitations. For concentration around *c* = 4.0 mg/ml, dense phase formation becomes much more evident with the appearance of a droplet (**Figure 2a**). At these concentrations, the chain *R*_*g*_, averaged over the whole system, has a rather wide distribution and lies between the *R*_*g*_ in the dense phase and that of a single chain (**Figure 2b**). As the concentration was increased further, the cluster size increased until it saturated at 500, signifying that all chains formed a single condensate. Correspondingly, *R*_*g*_ continued to increase, approaching the value of the dense bulk phase, 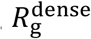. Note that the small discrepancy between the *R*_g_ measured in our condensate simulations and 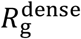 likely originates from slightly collapsed chains located at the condensate interface^44^. These findings reveal a gradual increase in average chain size as the system transitions from the dilute phase to the dense phase.

To substantiate the generality of these conclusions, we analyzed all EKVs (**Figures S6, S7**) and normalized the *R*_*g*_ and concentrations (**Figure 2c**). We normalized the *R*_*g*_ to span between the value for a single chain, 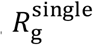, and the value of a chain in the dense bulk phase, 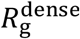. We normalized the concentrations by the *c*_*sat*_ values. We note that the *c*_*sat*_ values for EKV10 to EKV15 were extrapolated based on the data from sequences with lower nSCD, owing to the scarcity of chains in the dilute phase during slab co-existence simulations (**Figure S2b**). These extrapolated *c*_*sat*_ values did not affect the overall trend, as all the corresponding *R*_*g*_ values were similar to 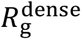. For EKVs with a low nSCD value (i.e., EKV2 to EKV7), *R*_*g*_ closely approximated 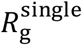 at concentrations below or slightly above the saturation concentration. At higher concentrations, where a distinct condensate formed, *R*_*g*_ increased until it almost reached 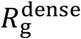. For the EKVs with higher nSCD values, the smallest concentration that we explored in our simulations is much larger than its saturation concentration, hence we did not observe a *R*_*g*_ value close to that of a single chain. However, with an increase in concentration, we still noted a rise in *R*_*g*_ until it plateaued at a value marginally smaller than 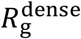. The normalized *R*_*g*_ values for all EKVs collapse onto a single sigmoidal curve (*R*^2^ = 0.95), exhibiting a uniform trend of increase from the dilute to the dense phase. This convergence demonstrates that the gradual increase in *R*_*g*_ from the dilute phase to the dense phase is a universal attribute across all EKVs.

Having established that for EKVs bearing zero net charge with diverse charge patterns, *R*_*g*_ progressively increases from the dilute phase to the dense phase, we next sought to verify this conclusion for natural IDPs. We selected four proteins, previously demonstrated to undergo LLPS *in vitro*, namely the low-complexity (LC) domain of FUS^45^, the disordered C-terminal domain of TDP-43^46^, the LC domain of hnRNPA2^47^ and the N-terminal disordered RGG domain of LAF-1 (LAF-1 RGG)^48^. According to previous research on phase behavior of these IDPs^49^, we conducted simulations at *T* = 300 K for the first three proteins, and at *T* = 260 K for LAF-1 RGG. We maintained a constant total monomer count of approximately 25,000 while varying the box volume to explore a range of concentrations from 0.2 mg/ml to 400 mg/ml. For all four sequences, the average *R*_*g*_ of the entire system initially remained nearly constant when the proteins remained dispersed in solution, and the average cluster size was close to one. As the dense phase formed (indicated by the growing cluster size shown in **Figure 3**), the average *R*_*g*_ gradually increased, eventually reaching the *R*_*g*_ that is characteristic of the bulk dense phase (**Figure 3, Figure S8**). The progression of *R*_*g*_ with increasing concentration is similar to those observed for the EKVs, with *R*_*g*_ gradually increasing alongside cluster size as the system evolves from the dilute phase into the dense phase. These findings suggest that during the phase separation process, IDPs undergo a transition from a dilute state to an oligomeric state, and ultimately to a dense state, accompanied by a gradual chain expansion.

**Figure 3.**
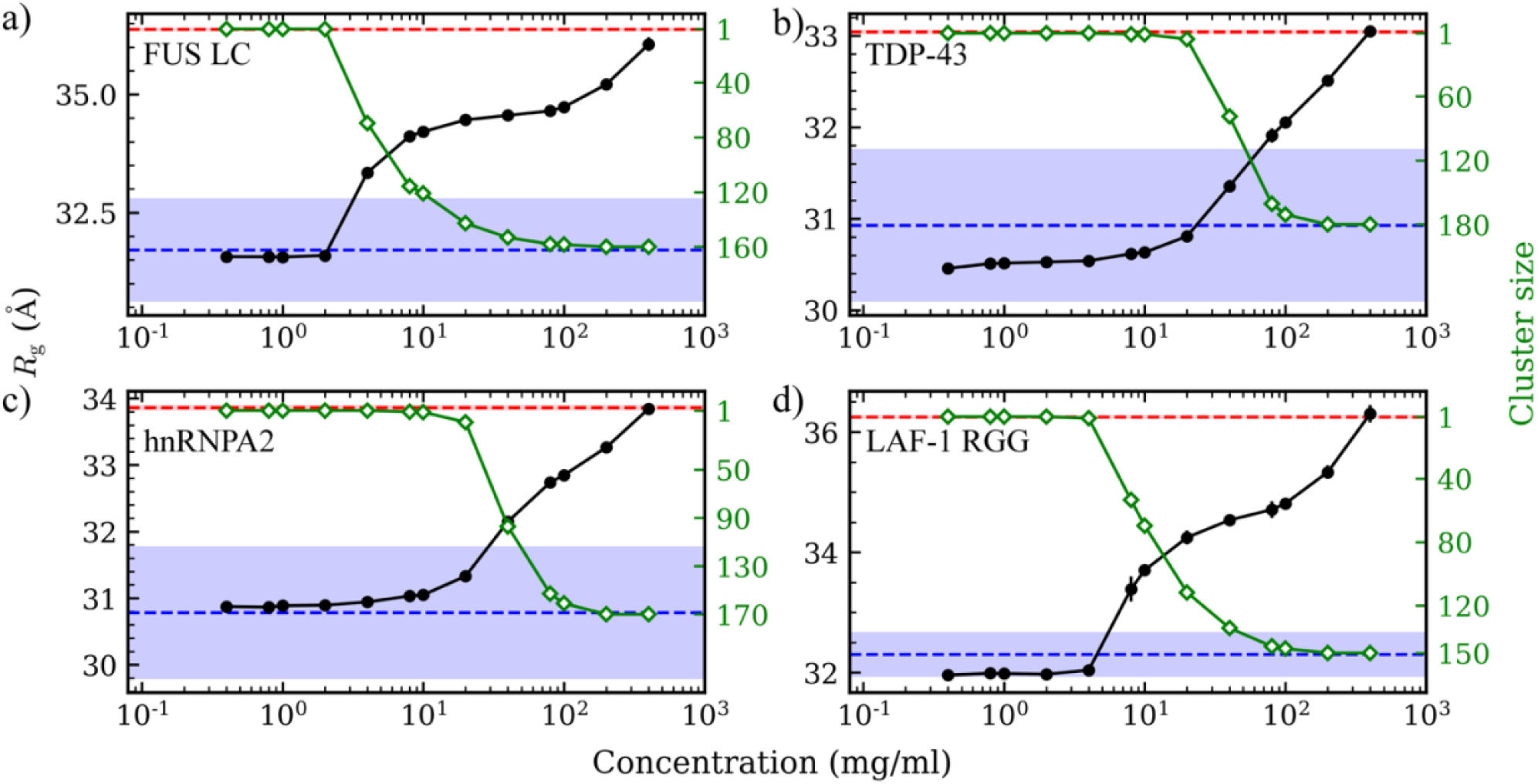
Concentration scan for the disordered domains of natural proteins: (a) FUS LC, (b) TDP-43, (c) hnRNPA2, and (d) LAF-1 RGG. *R*_g_ (black, left *y*-axis) and cluster size (green, right *y*-axis) as functions of concentrations. The red and blue dashed horizontal lines represent the *R*_g_ in the bulk dense phase and of a single chain, respectively.

Upon condensation, an interfacial region emerges between the dilute and dense phases, which likely plays an important role in the functionality and stability of MLOs^50–52^. Previously, our group elucidated the conformation of homopolymer chains and select IDPs at the interface of condensate droplets, and here we conducted a similar analysis for EKVs^44^. To eliminate the effects of (local) curvature of the condensate interface in a droplet geometry, we performed slab simulations to study conformations at interfaces. Taking EKV5 as an illustrative example (data for the other sequences are presented in the SI), a dense phase was observed, with occasional appearances of several chains in the dilute phase (**Figure 4a**). Interface boundaries were determined by fitting the concentration profile relative to the distance from the condensate center-of-mass to monomer in the *z* direction (*d*_zCOM_) using a hyperbolic tangent function^53^:

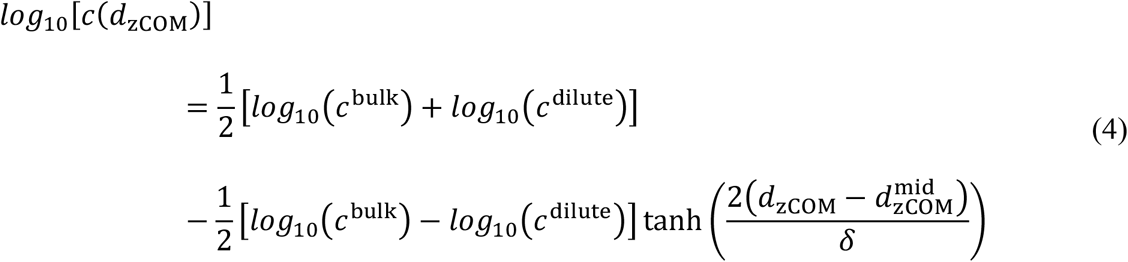

where *c*(*d*_zCOM_) is the concentration profile along the *z* direction, *c*^bulk^ and *c*^dilute^ are the concentrations of the bulk phase and dilute phase, respectively, 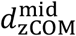 is the midpoint of the hyperbolic tangent function and δ is the width of the interface. Consequently, the interface boundaries are defined as 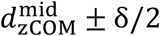.

**Figure 4.**
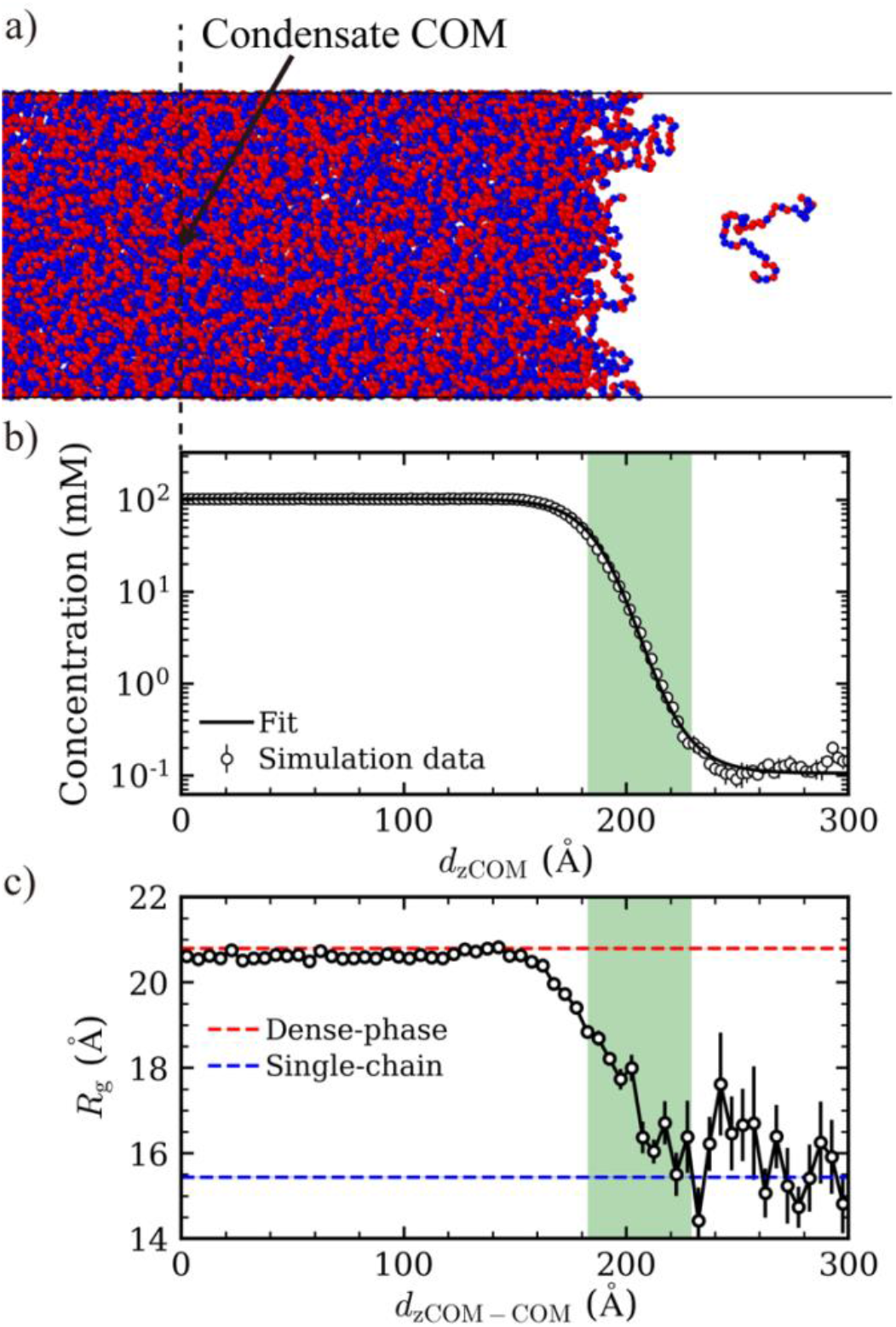
Analysis of EKV5 conformations at the interface *via* slab simulation. (a) Snapshot depicting one side of the interface of a condensate from the slab simulation. (b) Concentration profiles with respect to the distance from the condensate center-of-mass in the *z*-direction, *D* _zCOM_. The solid line is the fitted curve for the simulation data (symbols). (c) Average *R*_g_ with respect to distance from the condensate center-of-mass to chain’s center-of-mass in the *z* direction, *D* _zCOM−COM_. The red and blue dashed horizontal lines represent the *R*_g_ in the bulk dense phase and of a single chain, respectively.

Upon identifying the interfacial region (**Figure 4b**), we analyzed the local *R*_*g*_ with respect to the distance along the *z*-direction between a chain’s center-of-mass and the center-of-mass of the condensate, *D* _zCOM−COM._ In the dense region of the slab, *R*_*g*_ overlapped with the value obtained in the bulk dense phase, 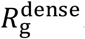. Within the interface region, *R*_*g*_ decreased and remained smaller than those observed for the chains within the bulk phase. Within the dilute phase, *R*_*g*_ exhibited pronounced fluctuations around the size of a single chain, 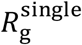. These fluctuations can be attributed to the limited statistical data available due to the low concentration of chains in the coexisting dilute phase. Importantly, all EKVs exhibit similar qualitative behavior, with conformations in the interfacial region being more compact than those in the dense interior and less compact than the dilute phase (**Figures S9, S10**).

In this simulation study, we investigated the conformational properties of proteins during phase separation by simulating model sequences composed of an equal number of oppositely charged monomers arranged in different patterns (referred to as EKVs) and naturally occurring disordered proteins. Despite extensive previous research on phase separation^1,2^, the description of conformational transitions underlying the condensation process remains largely qualitative. Due to the heterogeneous and dynamic nature of protein ensembles, it is difficult to tackle this question experimentally, which makes the simulation approaches used here an attractive avenue for gaining a comprehensive and quantitative understanding of sequence-dependent changes in conformational properties.

We decipher the self-assembly process itself by conducting simulations with increasing protein concentration. As expected, proteins initially stay as individual molecules in the dilute phase at low concentrations but then start to assemble into larger clusters at concentrations only above their saturation concentration. The cluster size grows with increasing protein concentration and eventually a single protein droplet forms with all proteins incorporated in it at very high concentrations. Importantly, ensemble conformations progressively shift from single-chain to bulk dense-phase and follow a sequence-independent universal behavior as a function of protein concentration normalized by the saturation concentration.

Even though the EKVs and natural proteins exhibit pronounced sequence-dependent conformational characteristics in the dilute phase (**Figure S11a**)^35^, the conformation within the dense phase demonstrates only a very weak sequence dependence (**Figure 5**). In fact, the relationship between the protein’s radius tion (*R*_*g*_) and end-to-end distance (*R*_e_) within the condensates is consistent with theoretical predictions for an ideal chain 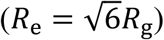 and their intramolecular distance (*R*_*ij*_) closely mirrors the scaling expected for an ideal chain (**Figure S11b**). These analyses strongly suggest that the protein conformations within the dense phase are akin to that of an ideal chain, which is characterized by being entropy-driven and not strongly influenced by the protein sequence or dilute-phase conformational properties^43^. We note that our results are consistent with the limited experimental data available in the literature^7,30,32,54^.

**Figure 5.**
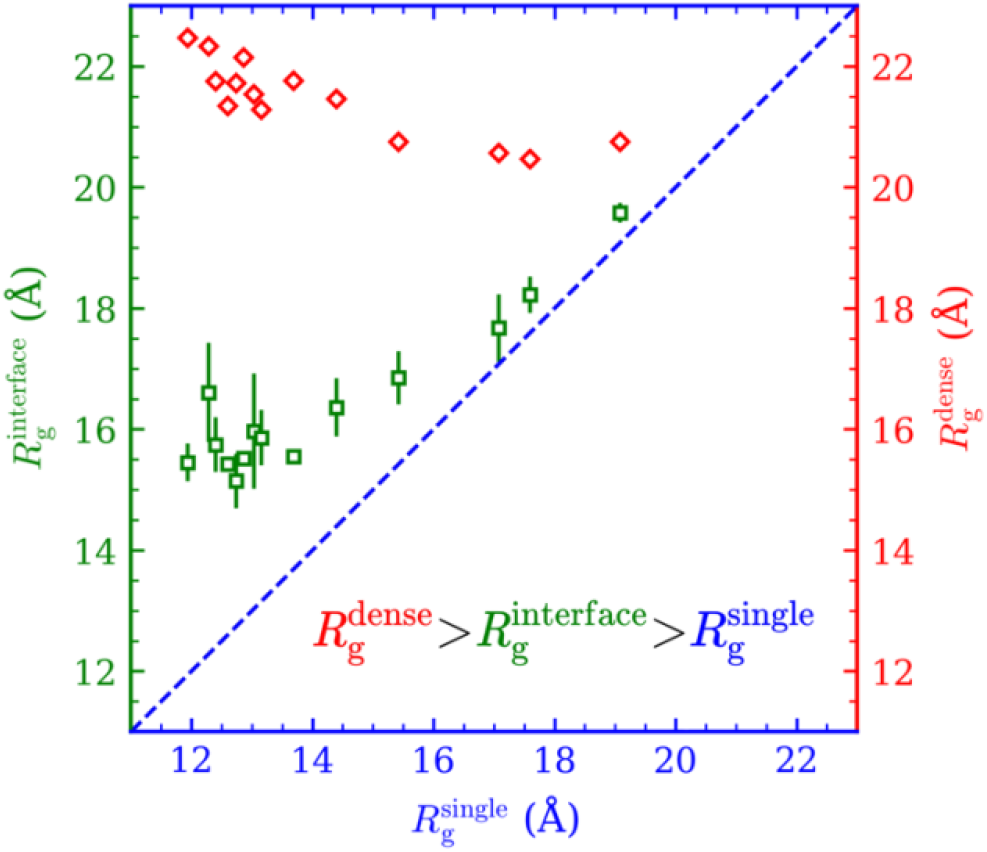
Correlation between 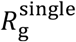 and 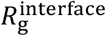 (green squares, left *y*-axis) or 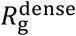 (red diamonds, right *y*-axis) for all investigated EKVs.

For conformational characteristics of proteins at the droplet interface, we do observe a significant dependence on the protein sequence, and hence their dilute-phase properties (**Figure 5**). Importantly, the protein chains at the interface are always more compact than the chains in the dense phase but remain more extended as compared to those in the dilute phase, independent of their sequence.

The results reported here uncover the general conformational characteristics of proteins during the phase separation process and highlight their distinct preferences as they move from the dilute phase to the dense phase including their interfacial behavior. These results should help understand the functional and pathological roles of conformational transitions associated with the formation of biomolecular condensates.

## Supporting Information

The details of the model and simulations, method of nSCD calculation, method for cluster analysis, amino acid code for the model and natural proteins, dense phase and saturation concentrations, probability distribution of *κ*^2^, cluster size distributions, effect of concentration scan on probability distribution of *R*_*g*_, cluster sizes, and average *R*_*g*_, concentration profile and *R*_*g*_ with respect to the distance from the condensate’s center of mass for the EKVs, interresidue distance analysis for the EKVs.

## Acknowledgments

This material is based on the research supported by the National Institute of General Medical Science (NIGMS) of the National Institutes of Health under the grant R01GM136917 and the Welch Foundation under the grant A-2113-20220331. The data on TDP-43 disordered domain were generated as part of a project supported by NINDS and NIA grant R01NS116176. A.N. acknowledges funding by the Deutsche Forschungsgemeinschaft (DFG, German Research Foundation) through Project 470113688. Y.C.K. is supported by the Office of Naval Research via the U.S. Naval Research base program. We gratefully acknowledge the computational resources provided by the Texas A&M High Performance Research Computing (HPRC) to complete this work.

