## Supporting Information for "Sequence-Dependent Conformational Transitions of Disordered Proteins During Condensation"

### Simulation details

We have used a coarse-grained model for the intrinsically disordered proteins (IDPs), in which each monomer (residue) is represented by a single bead. Beads are bonded *via* a harmonic spring:

$$U_{\mathrm{b}}(r) = \frac{k_{\mathrm{b}}}{2}(r - r_0)^2, \tag{1}$$

where $r$ is the distance between two bonded beads, $k_{\mathrm{b}}$= 20 $\mathrm{kcal}/(\mathrm{mol}\ \text{Å}^2)$ is the spring constant, and $r_0$ = 3.8 Å is the equilibrium bond length. The van der Waals interactions between nonbonded beads were modeled using a modified Lennard-Jones (LJ) potential[1]

$$U_{\mathrm{vdW}}(r) = \begin{cases} U_{\mathrm{LJ}}(r) + (1-\lambda)\varepsilon, & r \leq 2^{1/6}\sigma \\ \lambda U_{\mathrm{LJ}}(r), & otherwise \end{cases}, \tag{2}$$

where $U_{\mathrm{LJ}}$ is the standard LJ potential

$$U_{\mathrm{LJ}}(r) = 4\varepsilon\left[\left(\frac{\sigma}{r}\right)^{12} - \left(\frac{\sigma}{r}\right)^{6}\right], \tag{3}$$

with average hydropathy $\lambda = (\lambda_i + \lambda_j)/2$, average diameter $\sigma = (\sigma_i + \sigma_j)/2$ for residues *i* and *j*. The hydropathy values for E and K were $\lambda_{\mathrm{E}} = 0.460$ and $\lambda_{\mathrm{K}} = 0.514$, and diameters of the E and K residues were $\sigma_{\mathrm{E}} = 5.92$Å and $\sigma_{\mathrm{K}} = 6.36$Å, respectively[2,3]. For natural proteins, $\lambda_i$ values were set according to the HPS-Urry model[4]. The interaction strength was fixed to $\varepsilon = 0.2\,\mathrm{kcal/mol}$ in all simulations. For computational efficiency, the pair potential $U_{\mathrm{vdW}}$ and its associated forces were truncated to zero at a distance of 4 σ. The electrostatic interactions between nonbonded residues were modeled using a Coulombic potential with Debye–Hückel electrostatic screening[5]

$$U_e(r) = \frac{q_i q_j}{4\pi D \epsilon_0 r} e^{-r/\ell}, \quad (4)$$

where $D$ = 80 was the dielectric constant of the medium, $\epsilon_0$ was the permittivity of vacuum, and $\ell$=10 Å was the Debye screening length. The electrostatic potential and its forces were truncated to zero at a distance of 35 Å.

Single chain simulations were performed by placing one chain into a cubic box of edge length 160 Å with periodic boundary conditions in all directions (the simulation box was large enough to prevent unphysical self-interactions). Langevin dynamics simulations were performed at constant temperature ($T$ = 300 K for EKVs, FUS LC, TDP-43 and hnRNPA2, $T$ = 260 K for LAF-1 RGG). The friction coefficient was set as $\gamma_i = m_i/t_{damp}$, where $m_i$ is the mass of residues and $t_{\mathrm{damp}}$ = 1000 fs. Simulations were performed for 1.0 μs with a time step of 10 fs using HOOMD-blue[6] (ver. 2.9.3) with features extended using azplugins[7] (ver. 0.10.2). The simulation trajectories were saved every 1000 fs.

Dense phase simulations were performed by placing chains into a cubic box at a constant pressure of $P$ = 0 atm and equilibrating the systems for 0.5 μs. After the chains achieved their preferred dense-phase concentration, the simulations were performed using Langevin dynamics at constant volume for 1.0 μs. All the simulations were done using $t_{\mathrm{damp}}$ = 1000 ps and at fixed temperatures. Other simulation settings were also the same as the single chain simulations.

Concentration scan simulations were performed by placing a fixed number of chains ($N$ = 500) into cubic boxes of variable edge length, and then using Langevin dynamics ($t_{\mathrm{damp}}$ = 1000 ps) at specific constant volumes for 2 μs. Other simulation settings were the same as the single chain simulations.

Slab simulations were conducted by first preparing the dense phase in a cubic box of edge length 150 Å with periodic boundary conditions in all directions for 100 ns. Then, the z-dimension of the box was extended to 1200 Å, and simulations were performed for 3 μs using Langevin dynamics ($t_{\text{damp}}$ = 1000 ps). Other simulation settings were the same as the single chain simulations.

**Normalized sequence charge decoration parameter (nSCD):**

$$\text{SCD} = \frac{1}{N}\sum_{i=2}^{N}\sum_{j=1}^{i-1} q_i q_j (i-j)^{1/2}, \tag{5}$$

$$\text{nSCD} = \frac{\text{SCD}-\text{SCD}_{\text{max}}}{\text{SCD}_{\text{min}}-\text{SCD}_{\text{max}}}, \tag{6}$$

where $q_i$ and $q_j$ are the charges of residues *i* and *j*, respectively. $\text{SCD}_{\text{max}}$ is the SCD value for the perfectly alternating sequence and $\text{SCD}_{\text{min}}$ is for charge segregated sequence.

**Natural protein sequences used in this work**

FUS LC (length = 163)

MASNDYTQQATQSYGAYPTQPGQGYSQQSSQPYGQQSYSGYSQSTDTSGYGQSSYSSY
GQSQNTGYGTQSTPQGYGSTGGYGSSQSSQSSYGQQSSYPGYGQQPAPSSTSGSYGSSS
QSSSYGQPQSGSYSQQPSYGGQQQSYGQQQSYNPPQGYGQQNQYNS

TDP-43 (length = 141)

GRFGGNPGGFGNQGGFGNSRGGGAGLGNNQGSNMGGGMNFGAFSINPAMMAAAQAA
LQSSWGMMGMLASQQNQSGPSGNNQNQGNMQREPNQAFGSGNNSYSGSNSGAAIGW
GSASNAGSGSGFNGGFGSSMDSKSSGWGM

hnRNPA2 (length = 152)

GRGGNFGFGDSRGGGGNFGPGPGSNFRGGSDGYGSGRGFGDGYNGYGGGPGGGNFGG
SPGYGGGRGGYGGGGPGYGNQGGGYGGGYDNYGGGNYGSGNYNDFGNYNQQPSNY
GPMKSGNFGGSRNMGGPYGGGNYGPGGSGGSGGYGGRSRY

LAF-1 RGG (length = 168)

MESNQSNNGGSGNAALNRGGRYVPPHLRGGDGGAAAAASAGGDDRRGGAGGGGYRR
GGGNSGGGGGGGYDRGYNDNRDDRDNRGGSGGYGRDRNYEDRGYNGGGGGGGNRG
YNNNRGGGGGGYNRQDRGDGGSSNFSRGGYNNRDEGSDNRGSGRSYNNDRRDNGGD
G

**Cluster analysis**

We defined two chains as part of the same cluster if the distance between their monomers was less than 1.5σ (equivalent to 9.54 Å). Subsequently, we calculated the probability $P(N_C)$ of a chain being part of a cluster of size $N_C$. Following this, by summing the products of $P(N_C)$ and $N_C$ for all clusters, we derived the average cluster size.

**Supplementary Figures**

| EKV | | nSCD |
|---|---|---|
| 1 | EKEKEKEKEKEKEKEKEKEKEKEKEKEKEKEKEKEKEKEKEKEKEKEKEK | 0.000 |
| 2 | KEKKKEKEEEKKEKEKKEEKEKEEKKEEKKEEEKKKEEEKKEKKKEEEEK | 0.017 |
| 3 | KEKKKEKEEEKEEEKKEEKEEEKEEKKKKKEEEKKKEKEEEKEKKKEKEK | 0.033 |
| 4 | KKEEKEKEEEKEEEEEEKKKKEEKEKEKKKEKKEEEKKKEEKKKKEEEKK | 0.059 |
| 5 | KKEEKEKKEEEEEKEEEKEEKKKEEEEKKKKKKKKEKKEKEEEKEKKEEK | 0.067 |
| 6 | EEKKEKEEEEEEKEKEEEEEEKEEKEKKKKKKKKEKEKKKEKKKEKEEKK | 0.200 |
| 7 | EEEKEKEKEKEEEEEEEEEEEKEEKEKEKKKKKKEEKKKEKKKKKEKKKK | 0.333 |
| 8 | EKEKEEEEEEKEEEEEEEEKEEEKKEKEKKKKKKEEEKEKKKKKKKKKKK | 0.400 |
| 9 | EEEEEEEEEEKEEEEEKEEKKKEKKEKEEEKKKKEEKEKKKKKKKKKKKK | 0.468 |
| 10 | EEEEEEEEEEKEEEEEEEEKKKEKKKKEEEKKKKEKKEKKEKKKKKKKKK | 0.500 |
| 11 | EEEEEEEEEEEEEEEEKEEKKEKEEKKKEKKKEKEKKKKEKKKKKKKKKK | 0.601 |
| 12 | EEEEEEEEEEEEEEEEKEEEKEKEKEKEKKKKKKKKEKKKKKKKKEKKKK | 0.667 |
| 13 | EEEEEEEEEEEEEEEKEEEEEKKKEEEEKKKKKKKKKKEKKKKKKKKKKK | 0.734 |
| 14 | EEEEEEEEEEEEEEEEEEEEEKEKEKEKEKKKKKKKKKKKKKKKKKKKKK | 0.902 |
| 15 | EEEEEEEEEEEEEEEEEEEEEEEEEKKKKKKKKKKKKKKKKKKKKKKKKK | 1.000 |

**Figure S1.** Selected E-K variants (EKVs) for chain length N=50 with their identifying number and normalized SCD parameter.

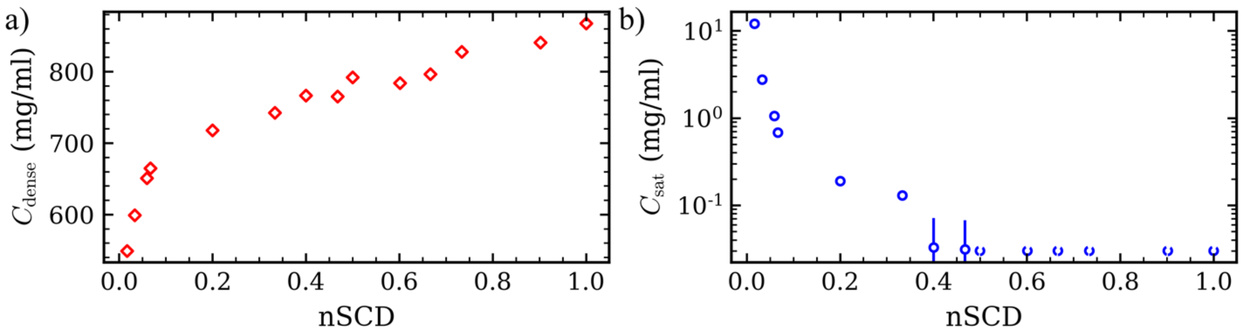


**Figure S2.** (a) Dense phase concentrations for EKVs as a function of nSCD. (b) Saturation concentrations for EKVs as a function of nSCD. The open symbols represent the extrapolated concentrations for EKV10 to EKV15.

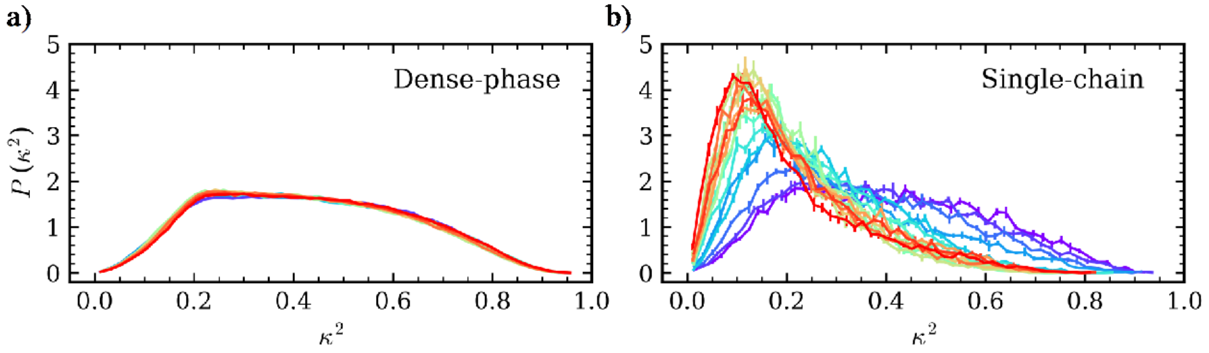


**Figure S3.** Probability distribution of relative shape anisotropy $\kappa^2$ for EKVs in the (a) dense phase and as (b) a single chain. Line colors ranging from purple to red, indicate increasing nSCD.

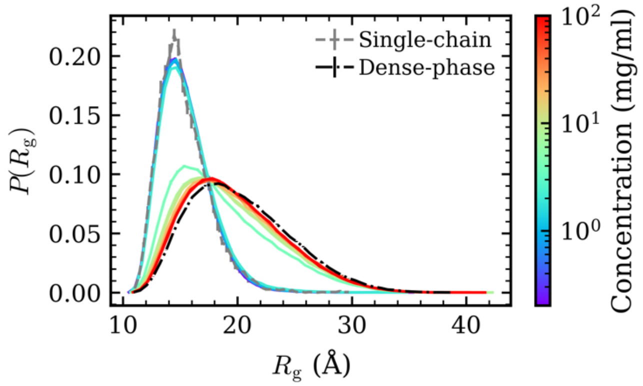


**Figure S4.** Probability distribution of $R_g$ for EKV5. Line colors ranging from purple to red, indicate increasing concentrations from 0.2 mg/ml to 100 mg/ml. The gray dashed line represents the $R_g$ distribution for a single chain and the black dashed line represents the distribution within the dense phase.

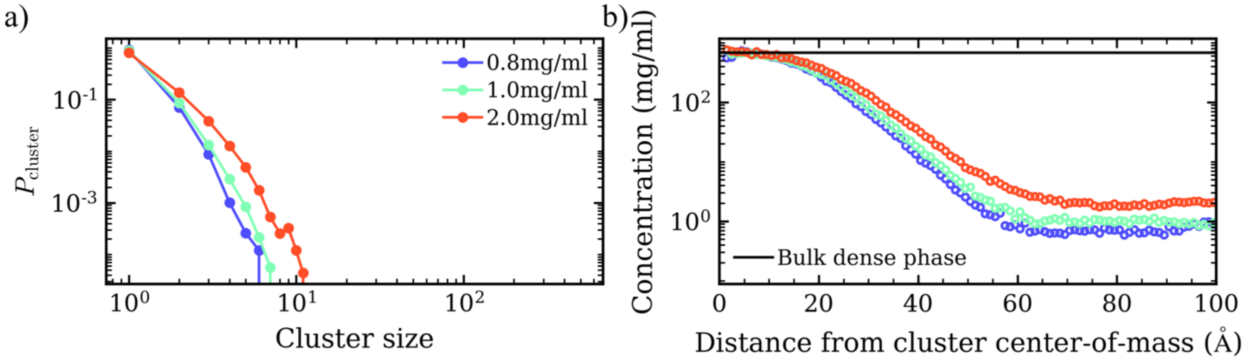


**Figure S5.** (a) Probability distribution of cluster size for EKV5 at the concentrations of 0.8 mg/ml, 1.0 mg/ml, and 2.0 mg/ml. (b) Radial density profile of monomers with respect to the distance from the center-of-mass of the largest cluster.

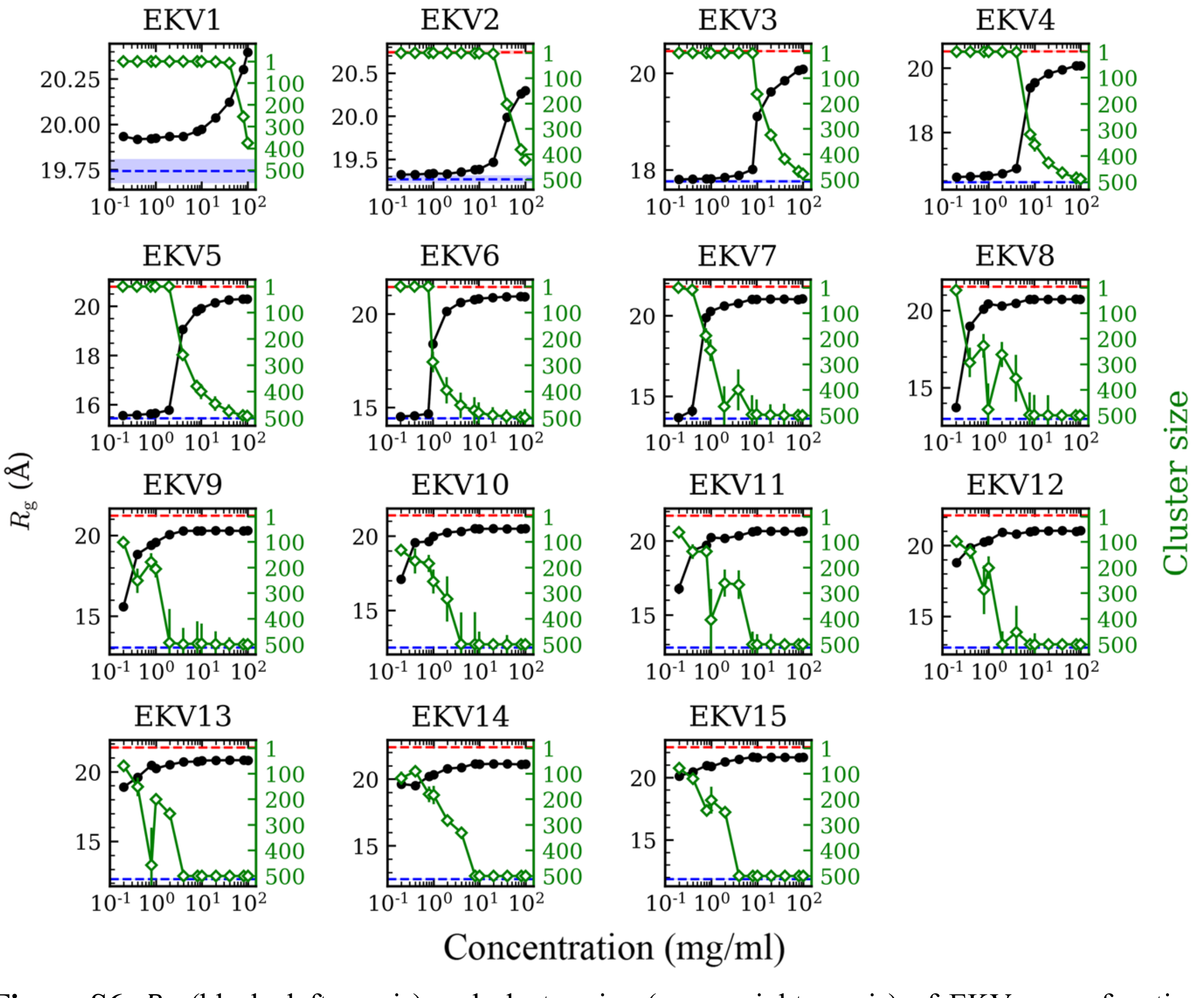


**Figure S6.** $R_g$ (black, left *y*-axis) and cluster size (green, right *y*-axis) of EKVs as a function of concentration. The red and blue dashed horizontal lines represent the $R_g$ in the bulk dense phase and of a single chain, respectively.

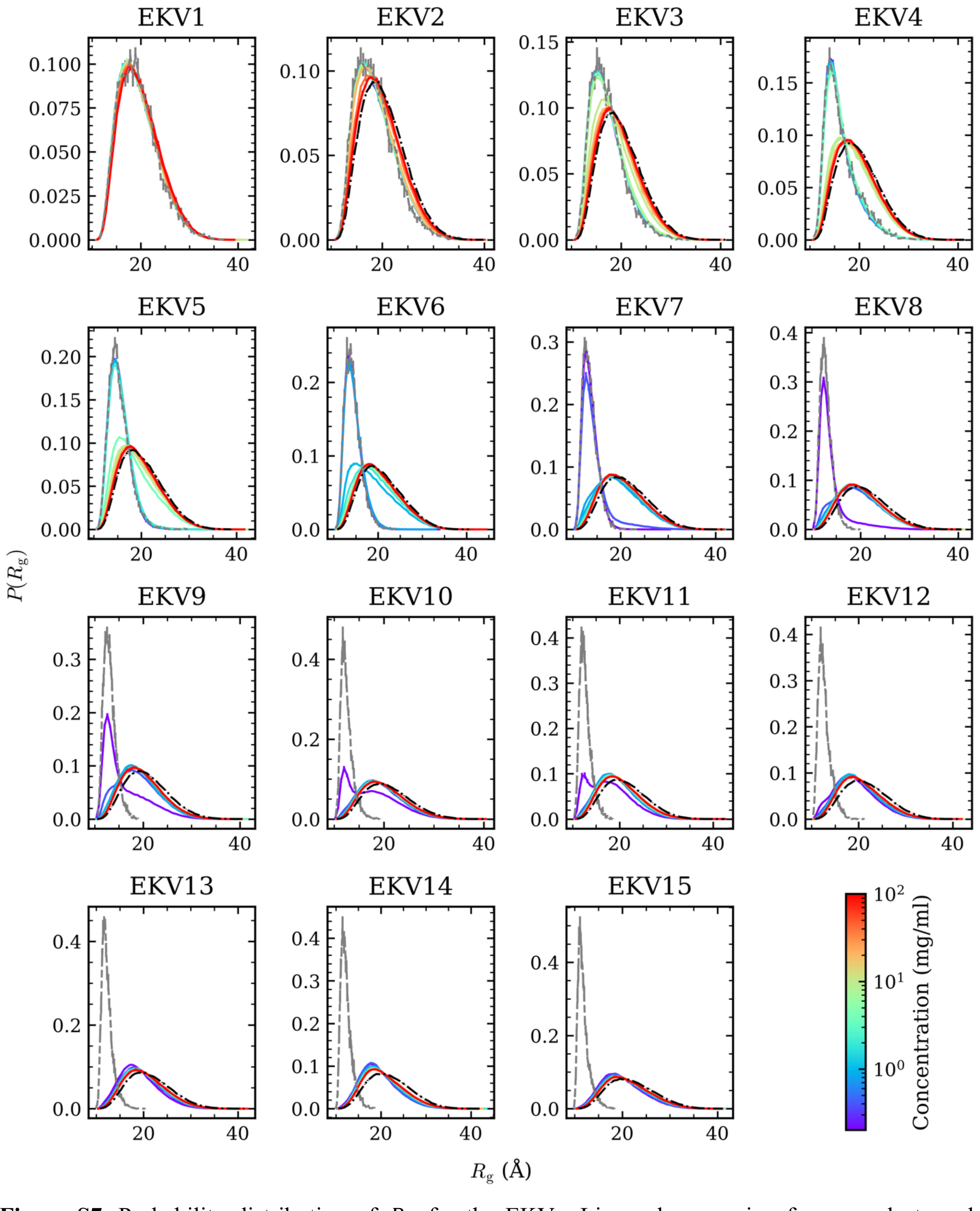


**Figure S7.** Probability distribution of $R_g$ for the EKVs. Line colors ranging from purple to red, indicate increasing concentrations. The gray dashed line represents the $R_g$ distribution for a single chain and the black dashed line represents the distribution within the dense phase.

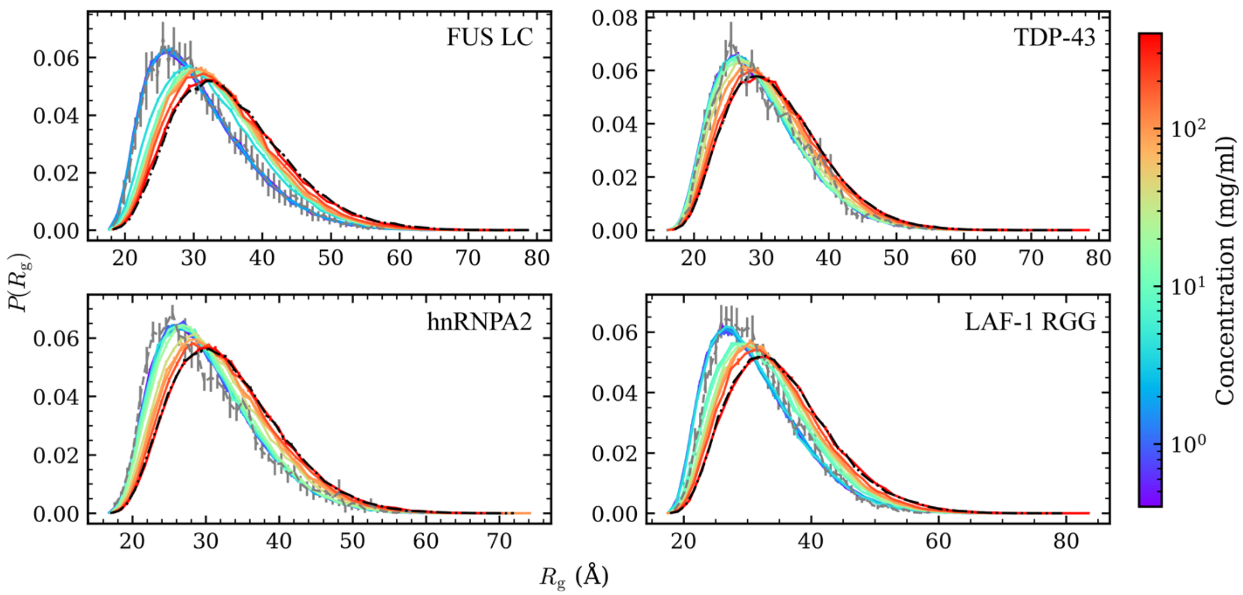


**Figure S8.** Probability distribution of $R_g$ for the disordered domains of natural proteins: FUS LC, TDP-43, hnRNPA2, and LAF-1 RGG. Line colors ranging from purple to red, indicate increasing concentrations for 0.2 mg/ml to 400 mg/ml. The gray dashed line represents the $R_g$ distribution for a single chain and the black dashed line represents the distribution within the dense phase.

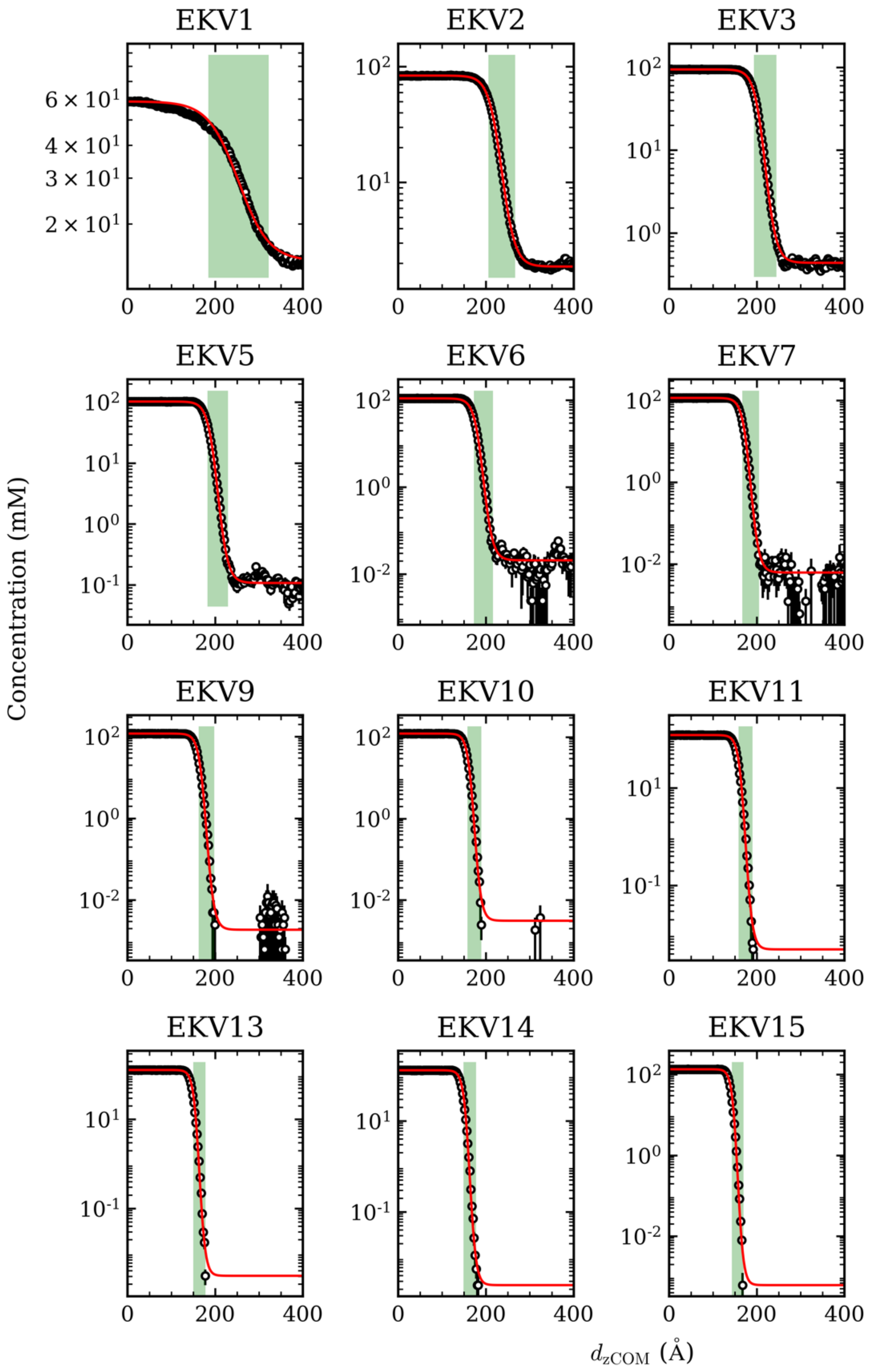


**Figure S9.** Concentration profiles with respect to the distance from the condensate center-of-mass in the *z* direction. The red line is the fitted curve for the simulation data (symbols). The green shaded area indicates the interface region.

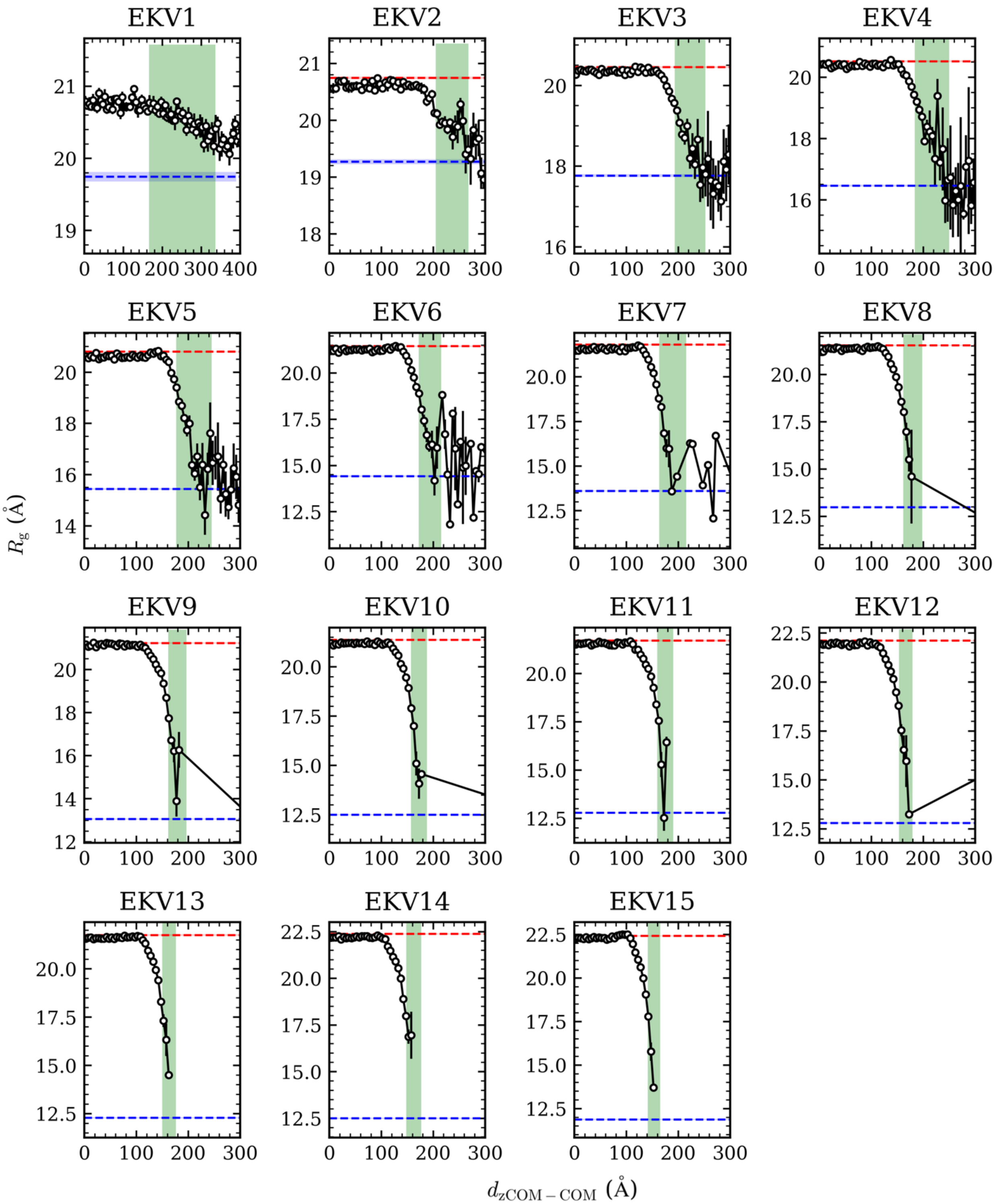


**Figure S10.** Interface analysis for the EKVs *via* slab simulations. Average $R_g$ with respect to distance from the condensate center-of-mass to chain's center-of-mass in the $z$ direction, $d_{zCOM-COM}$. The red and blue dashed horizontal lines represent the $R_g$ in the bulk dense phase and of a single chain, respectively.

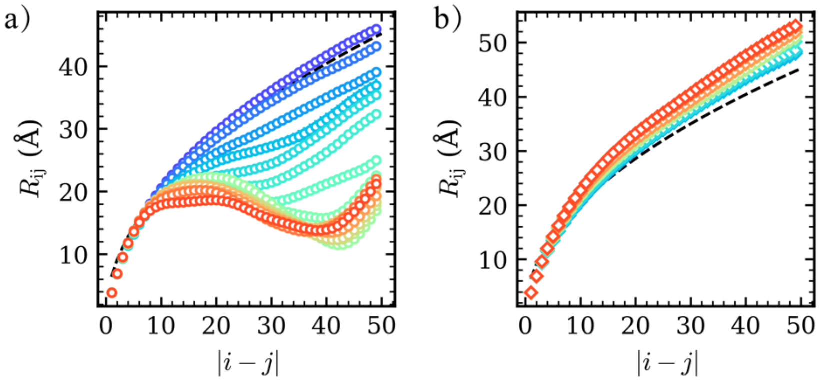


**Figure S11.** Interresidue distance $R_{\mathrm{ij}}$ as a function of residue separation in the chain $|i-j|$ for the EKVs in the (a) single-state and the (b) dense phase. The dashed line corresponds to the ideal chain scaling $R_{\mathrm{ij}} = b|i-j|^{1/2}$, where $b = 6.39$Å was fitted for EKV1 using the theoretically expected end-to-end distance $R_e^2 = Nb^2$ of an ideal chain with $N = 50$. The symbol color, ranging from purple to red, indicates increasing nSCD.

(7) https://github.com/mphowardlab/azplugins
